# Construction of a novel 3D urinary bladder mucosa model and its application in toxicity assessment of arsenicals

**DOI:** 10.1101/2025.08.19.670771

**Authors:** Runjie Guo, Min Gi, Tohru Kiyono, Arpamas Vachiraarunwong, Shugo Suzuki, Masaki Fujioka, Guiyu Qiu, Kwanchanok Praseatsook, Xiaoli Xie, Hideki Wanibuchi

**Affiliations:** Department of Environmental Risk Assessment, Osaka Metropolitan University Graduate School of Medicine, Osaka, Japan; Project for Prevention of HPV‑Related Cancer, Division of Collaborative Research and Development, Exploratory Oncology Research and Clinical Trial Center, National Cancer Center, Chiba, Japan; Department of Molecular Pathology, Osaka Metropolitan University Graduate School of Medicine, Osaka, Japan; Department of Toxicology, School of Public Health, Southern Medical University (Guangdong Provincial Key Laboratory of Tropical Disease Research), Guangzhou, China

**Keywords:** 3D urinary bladder mucosa model, Arsenic, Toxicity assessment, Urothelial cell marker, γ-H2AX

## Abstract

The urinary bladder is a major target organ for environmental toxicants, including arsenic. The objects of this study were two-fold. First, we constructed a novel 3D urinary bladder mucosa model that incorporated an overlying epithelium and a supporting subepithelial layer, referred to as the 3D-UBMM. Primary human bladder urothelial and fibroblast cells were immortalized by introducing the human CDK4^R24C^ and TERT genes. The subsequent construction of the 3D-UBMM involved incorporating the immortalized fibroblast cells into a collagen raft, while the immortalized urothelial cells were cultured at the air-liquid interface of the raft. This 3D-UBMM closely resembles the human urinary bladder epithelium in terms of morphology and marker protein expression, including uroplakin 1b, P63, and cytokeratin 5. Second, using the 3D-UBMM we investigated the cytotoxicity of sodium arsenite (iAs^III^) and dimethylarsenic acid (DMA^V^). Exposure to iAs^III^ and DMA^V^ resulted in increased urothelial cell necrosis and increased γ-H2AX-positive cells along with a reduction of P63-positive cells, and each of these effects was induced in a dose-response manner. These findings affirm that this novel 3D-UBMM closely resembles the human urinary bladder epithelial layer, offering a practical *in vitro* model for the evaluation of the toxicity of arsenic and other bladder carcinogens and the role of cancer-related genes in bladder carcinogenicity. In addition, by identifying mechanisms of carcinogenesis this model will aid in hazard identification and risk assessment of potential bladder carcinogens.

## 1. Introduction

The urinary bladder is responsible for storing and voiding urine (Atala, 2011). The urinary bladder wall consists of four distinct layers of tissue structure: the mucosa, the submucosa, the muscularis, and the parietal peritoneum. The mucosal layer, also known as the urothelium, lines the urinary bladder and is composed of three layers of specialized epithelial cells: an apical layer of umbrella-shaped cells, a middle intermediate layer of epithelial cells that can be several cells thick, and a basal stem cell layer. The stem cell layer supports regeneration of the upper epithelial cells in response to injury and also provides injury induced signals to activate underlying stromal fibroblasts (Shin et al., 2011). The submucosa contains urinary bladder fibroblasts, which provide both physical support and nutrition to the overlying epithelial cells. These fibroblasts engage in the synthesis of extracellular matrix and play a crucial role in the maturation and homeostasis of the urothelium (Shin et al., 2011; Wang et al., 2017). The main role of the mucosal layers is to function as an osmotic and chemical barrier, protecting the underlying tissues from potential damage caused by urine (Fry and Vahabi, 2016). Urine in the bladder accumulates the terminal products of the body’s metabolism, leading to frequent exposure of the urinary bladder epithelium to various harmful factors, including carcinogens and toxins. In addition, a wide range of pathogens cause urinary tract infections. Therefore, most bladder diseases are closely linked to damage to the bladder epithelium.

Generally, two-dimensional (2D) cell culture has been used to study the physiopathological mechanisms of bladder-related diseases. However, 2D cell culture cannot accurately mimic the *in vivo* condition of the urinary bladder epithelium. In our body, the urinary bladder epithelium is closely connected to the underlying connective tissue containing bladder fibroblast cells, and this unique microenvironment influences cell behavior, including proliferation, differentiation, and apoptosis (Du et al., 2021). In recent years, the widespread use of three-dimensional (3D) cell models of tumor and skin epithelium has also provided valuable insights into the study of bladder-related diseases (Gomez-Roman et al., 2017; Rikken et al., 2020). One of the main advantages of 3D models is that they provide a cellular model more in accordance with the physiological structure of the tissue. Moreover, the emergence of diseases is often a consequence of complex cellular interactions. Chen’s research indicated that inflammation in bladder fibroblast cells could impact overlying urothelial cells, thereby playing a pivotal role in bladder carcinogenesis (Chen et al., 2020). Therefore, when investigating the mechanisms of bladder-related diseases *in vitro*, 3D models are preferable to 2D cell culture systems. Inorganic arsenic is a known human carcinogen, linked to the development of urinary bladder, lung, and skin cancers. The International Agency for Research on Cancer (IARC) categorizes arsenic and inorganic arsenic compounds as Group 1 carcinogens (carcinogenic to humans) (IARC, 2012). Chronic arsenic exposure remains a significant public health issue in many countries. Human exposure to arsenic primarily occurs through drinking contaminated water, as well as through inhalation, food and skin contact (Chen et al., 2009). When arsenic-containing water is consumed, it is metabolized in the body into two forms, inorganic arsenic and organic arsenic, both of which are closely related to the incidence of urinary bladder cancer (Cohen et al., 2019). Inorganic arsenic has two main forms in the body, pentavalent (iAs^V^) and trivalent (iAs^III^). iAs^III^ is more rapidly absorbed than iAs^V^ and exhibits significant cytotoxicity even at low concentrations. Its quick uptake and high reactivity trigger cellular oxidative stress and inflammation in the urinary bladder epithelium, fostering DNA damage and irregular cell growth, potentially resulting in carcinogenesis.

The organic arsenical Dimethylarsenic acid (DMA^V^) is classified as a Group 2B compound, possibly carcinogenic in humans, by IARC (IARC, 2012). DMA^V^ is easily absorbed through the intestines, subsequently distributing to organs such as the kidneys, urinary bladder, and skin (Cohen et al., 2013). DMA^V^ is a urinary metabolite of various inorganic arsenic compounds including iAs^III^, and has been shown to be closely associated with the development of urinary bladder cancer in rats, mainly through cytotoxicity, oxidative stress, inflammation, and other pathways that lead to abnormal proliferation of urothelial cells (Cohen et al., 2002; Wei et al., 2002). However, Cohen et al. report that metabolism of DMA^V^ resulting in production of high levels of Trimethylarsine oxide (TMAO) is unique to rats (Cohen et al., 2006). They show that in rats administered DMA^V^ the levels of TMAO in the urine is markedly higher than in mice and humans. They also note that during the formation of TMAO the highly reactive arsenical DMA^III^ is formed and reviews by Cohen et al. state that the mode of action of DMA^V^-induced bladder carcinogenesis in the rat involves the generation of DMA^III^ (Cohen et al., 2007, 2006). Cohen et al. also report that in all tested animal species, including humans, DMA^V^ does not readily translocate into intact cells and consequently DMA^V^ metabolites, including DMA^III^, are produced at low levels (Cohen et al., 2006). However, Cohen et al. also report that in the rat there is higher cellular uptake and metabolism of DMA^V^, and the rat is the only species in which administration of DMA^V^ results in urinary concentrations of DMA^III^ that are cytotoxic to urothelial cells *in vitro* (Cohen et al., 2006). In agreement with these reports, Arnold et al. found that administration of DMA^V^ caused bladder cancer in rats, but did not induce bladder hyperplasia or bladder cancer in mice (Arnold et al., 2006). This suggests that in this study the low translocation of DMA^V^ into cells in mice resulted in low metabolism of DMA^V^ to DMA^III^, resulting in urinary concentrations of DMA^III^ that did not reach cytotoxic levels in DMA^V^-treated mice. In addition, Dodmane et al. reported that administration of DMA^III^ to wild-type mice resulted in urothelial cytotoxicity and regenerative proliferation (Dodmane et al., 2013). Overall, current data indicates that in mice and humans the cytotoxic potential of DMA^V^ is less than that of iAs^III^.

Despite extensive animal experiments to explore the mechanisms of organic and inorganic arsenic toxicity, there is only a limited understanding of the mechanisms of toxicity and carcinogenicity in the human urinary bladder. To address this gap, we utilized primary human bladder urothelial cells and fibroblast cells to construct a 3D urinary bladder mucosa model that incorporated an overlying epithelium and a supporting subepithelial layer, referred to as the 3D-UBMM, that closely resembles the characteristics of the human urinary bladder mucosa. We also tested the applicability of our 3D-UBMM by assessing the cytotoxicity of an arsenical with stronger *in vivo* cytotoxicity, iAs^III^, and an arsenical with weaker *in vivo* cytotoxicity, DMA^V^.

## 2. Materials and Methods

### 2.1. Chemicals

Sodium arsenite (iAs^III^, purity > 90%) was purchased from Sigma-Aldrich (St. Louis, MO, USA). DMA^V^[(CH_3_)_2_AsO(OH)] (purity > 99%) was purchased from Fluka (Steinheim, Germany).

### 2.2. Human urinary bladder tissue

Normal human urinary bladder tissue was obtained from an autopsy case at Osaka Metropolitan University Hospital, Osaka, Japan. The Ethical Committee of the Osaka Metropolitan University Graduate School of Medicine approved the use of this human specimen and clinical data (Approval number #2021-047) in accordance with the Declaration of Helsinki and the guidelines of the Osaka Metropolitan University Graduate School of Medicine.

### 2.3. Establishment of immortalized urinary bladder fibroblast and epithelial cell lines

Primary human urinary bladder fibroblast cells (HBladFB) were purchased from ATCC® (ATCC® PCS-420-013TM, United States), and primary human urinary bladder epithelial cells (male Caucasian, 19 years old) (HBladEC) were purchased from KURABO (KP-4109, Osaka, Japan). To immortalize the HBladFB and HBladEC cells, lentivirus vectors CSII-CMV-TERT and CSII-CMV-CDK4^R24C^, expressing human telomerase reverse transcriptase (TERT) and CDK4^R24C^ (mutant CDK4 an INK4a-resistant form of CDK4), were introduced into each cell line using methods described previously (Inagawa et al., 2014; Nishiwaki et al., 2020). The resulting immortalized cell lines were designated HBladFB-K4T and HBladEC-K4T, and expressed both TERT and CDK4^R24C^.

HBladEC-K4T has been confirmed to express As3MT by western blot analysis and to be negative for P53 by immunochemistry (data not shown). Whole-exome sequencing using the SureSelect V8-Post kit on the Illumina platform, conducted by Cell Innovator Inc. (Fukuoka, Japan), identified a single nucleotide substitution at chr17:7676154 within exon 4 of the TP53 gene, resulting in an amino acid change from proline to arginine. Functional prediction using the SIFT algorithm (https://sift.bii.a-star.edu.sg/index.html) classified this variant as “tolerated,” suggesting that it is unlikely to have a significant impact on TP53 protein function (data not shown).

In addition, although the arsenic metabolic capacity of HBladEC-K4T cells was not directly examined in the present study, our previous work demonstrated that HBladEC-T cells, which are immortalized with TERT alone, are capable of metabolizing various arsenicals, including iAs^III^ and DMA^V^, via pathways consistent with known arsenic metabolism (Vachiraarunwong et al., 2024). In preliminary experiments, HBladEC-T cells also exhibited stratification and urothelial differentiation; however, the multilayer structure was thinner than that observed in HBladEC-K4T cells. This difference is likely due to the absence of mutant CDK4 (K4) in HBladEC-T cells. Since K4 is primarily involved in enhancing proliferative capacity and is not known to influence arsenic metabolism, we expect that the arsenic-metabolizing capacity of HBladEC-K4T cells is comparable to that of HBladEC-T cells.

The HBladFB-K4T cells were cultured in DMEM culture medium (Nacalai Tesque Inc., Kyoto, Japan) supplemented with 2% FBS, 5 ng/ml basic FGF, 50 mg/ml ascorbic acid, 1 mg/ml hydrocortisone hemisuccinate, 5 mg/ml recombinant human insulin, and 100u/ml penicillin and 100 mg/ml streptomycin, referred to as HBladFB-K4T medium. The HBladEC-K4T cells were cultured in F medium (DMEM/Ham’s F-12 culture medium (Nacalai Tesque Inc., Kyoto, Japan) containing FBS, hEGF, adenine-HCl, insulin, hydrocortisone, cholera toxin, and penicillin-streptomycin) (Liu et al., 2012) supplemented with 10mM Y-27632 (Y), 5% (v/v) conditioned medium from Wnt-3A cells expressing human RSPO-1 and human Noggin (purchased from ATCC (CRL-2647)), 50 nM A-83-01 (A), and 50 nM DMH1 (D), collectively referred to as FYWRAD medium.

### 2.4. Evaluation of cytotoxicity of iAs^III^ and DMA^V^ in 2D culture

The cytotoxicity of iAs^III^ and DMA^V^ to HBladEC-K4T epithelial cells was investigated by evaluating cell viability. Stock solutions of 10 mM iAs^III^ and 5 M DMA^V^ were prepared in 0.5 M HEPES (pH 7.2, Thermo Fisher Scientific, USA) and subsequently diluted to the appropriate working concentrations using FYWRAD medium. HBladEC-K4T epithelial cells were seeded at a density of 5 × 10^3^ cells/well into 96-well plates and incubated for 24 h. Subsequently, the FYWRAD medium was replaced with FYWRAD medium containing different concentrations of iAs^III^ and DMA^V^, and the cells were incubated for 24 h. After incubation with the arsenicals, cell viability was determined by Cell Counting Kit-8 (CCK-8, Dojindo Laboratories, Kumamoto, Japan). The FYWRAD medium containing the iAs^III^ and DMA^V^ was removed and fresh medium was added to each well. After adding CCK-8 solution into each well, the 96-well plates were incubated at 37 ºC in an incubator with 5% CO_2_ for 4 h. The ratio of the optical density (OD) of treated wells to the mean absorbance of control wells was calculated to determine the percentage of cell viability. The LC_20_ and LC_50_ doses were separately determined for the iAs^III^ and DMA^V^ and calculated by nonlinear regression analysis using GraphPad Prism 8 (Okuno et al., 2019).

### 2.5. Construction of the 3D-UBMM

To establish the 3D-UBMM, we first performed a series of optimization experiments to determine the most suitable conditions for supporting epithelial differentiation and maintaining the 3D-UBMM integrity. Various combinations of HBladEC-K4T epithelial cells (1 × 10^5^, 1.5 × 10^5^, 3 × 10^5^, 5 × 10^5^) and HBladFB-K4T fibroblast cells (0, 1 × 10^5^, 3 × 10^5^, 5 × 10^5^, 1 × 10^6^) were tested, along with different ratios of HBladFB-K4T medium to FWR medium (F medium supplemented with WR, see Section 2.3) at 3:1, 1:1, and 1:3. These experiments showed that HBladEC-K4T cells failed to differentiate and maintain normal morphology in the absence of fibroblasts, indicating that matrix cellularity is critical for establishing a stable and functional 3D model. As differences in matrix cellularity can influence not only tissue homeostasis but also toxicological outcomes—such as the epithelial response to arsenic exposure—by altering the microenvironmental support, we fixed the optimized fibroblast density to maintain consistency across all toxicological evaluations. Based on these results, we selected the optimized condition of 5 × 10^5^ fibroblasts embedded in a collagen matrix, 1.5 × 10^5^ epithelial cells seeded on top, and a 3:1 mixture of HBladFB-K4T medium and FWR medium. We also compared the impact of submerged versus air-liquid interface culture conditions on epithelial differentiation. Despite testing various culture media, including artificial urine solution (#900489945, Isekyu Co. Ltd., Nagoya, Japan), under submerged conditions, HBladEC-K4T epithelial cells consistently formed only a monolayer and failed to differentiate into uroplakin-positive umbrella cells (data not shown). Although the underlying mechanism remains unclear, our findings suggest that the air-liquid interface is essential for promoting stratification and urothelial differentiation *in vitro*.

To construct the model under these optimized conditions, 5 ml of HBladFB-K4T medium containing 5×10^5^ HBladFB-K4T fibroblasts, 1 ml of 0.5% Collagen IPC-50 (AteloCell, Japan), and 0.05 ml of 1M HEPES (Gibco, USA) were mixed and poured into one well of a 6-well plate (Sumitomo Bakelite. Japan). The 6-well plate was then placed in a 37 °C incubator to allow the mixture to solidify. This solid fibroblast-containing material is referred to as a collagen raft.

After 4-5 days, the collagen rafts shrunk to about 10 mm in diameter and were transferred to Falcon Cell Culture Inserts with 0.4 μm pores and a 10.5 mm diameter (Corning, USA, #353493) in a 12-well plate. Then, 0.1 ml of FYWRAD medium containing 1.5 × 10^5^ HBladEC-K4T epithelial cells were seeded onto the surface of the collagen rafts. An additional 750 μl of HBladFB-K4T medium was added into the bottom of the well.

After 24 hours, the HBladFB-K4T medium in the insert was removed, and the medium in the bottom of the well was replaced with 750 μl of a 3:1 mix of HBladFB-K4T medium and FWR medium to maintain HBladEC-K4T cells at the air-liquid interface. After nine days the cells had achieved the 3D-UBMM configuration. On day 10, arsenicals were added into the 3D-UBMM to evaluate their cytotoxicity.

### 2.6. Evaluation of the cytotoxicity of iAs^III^ and DMA^V^ using the 3D-UBMM

To assess the cytotoxicities of iAs^III^ and DMA^V^, 100 μl of FYWRAD medium containing targeted concentrations of iAs^III^ or DMA^V^ was added into the insert containing the 3D-UBMM, and 750 μl of the 3:1 mix of HBladFB-K4T and FWR medium was added into the bottom of the wells, and the cells were incubated at 37 ºC in an incubator with 5% CO_2_. Twenty-four hours after treatment, the 3D-UBMM was fixed in 10% phosphate-buffered formalin for one day and then cut into 3 strips, embedded in paraffin, and processed for hematoxylin/eosin and immunohistochemical staining.

### 2.7. Immunohistochemical staining

Paraffin-embedded sections of the human urinary bladder (Section 2.2) and the 3D-UBMM were stained using the avidin-biotin-peroxidase complex (ABC) method. Briefly, the section was deparaffinized, dehydrated, and the antigen was retrieved by microwaving at 98°C for 20 min in 0.01 M citrate buffer (pH 6.0). Next, the endogenous peroxidase activity was blocked with 3% H_2_O_2_ in distilled water for 5 min, followed by blocking of non-specific binding with 10% goat or horse serum at room temperature for 20 min. Sections were then incubated with primary antibodies overnight at 4°C. The primary antibodies and dilutions were as follows: Uroplakin 1b (UPK1B; ab263454, Abcam, USA, mouse monoclonal, 1:200), P63 (ab735, Abcam, USA, rabbit monoclonal, 1:1000), cytokeratin-5 (CK5; ab53121, Abcam, USA, rabbit polyclonal, 1:1000), γ-H2AX (9718S, CST, USA, rabbit monoclonal, 1:500). Reactivity with the primary antibody was detected by incubating the sections with biotin-labeled goat anti-rabbit IgG or biotin-labeled horse anti-mouse IgG using the VECTASTAIN ABC kit (Vector Laboratories, Burlingame, CA, USA) and diaminobenzidine tetrahydrochloride. Tissue sections were counterstained with hematoxylin.

### 2.8. Statistical analysis

Data are reported as mean ± standard deviation (mean ± SD). Statistical analyses were performed using Prism 8 software (GraphPad Software, Inc., San Diego, CA, USA). One-way ANOVA followed Dunnett’s multiple comparison test was used to compare differences comparison between multiple groups. P values less than 0.05 were considered statistically significant.

## 3. Results

### 3.1. Establishment of immortalized primary HBladEC and HBladFB cell lines, and construction of the 3D-UBMM

Fig. 1A illustrates the procedure of immortalization of HBladEC epithelial cells and HBladFB fibroblasts to HBladEC-K4T epithelial cells and HBladFB-K4T fibroblasts by introducing TERT and CDK4^R24C^. Fig. 1B shows the immortalized HBladEC-K4T epithelial cells and HBladFB-K4T fibroblasts. A schematic illustration of the 3D-UBMM is shown in Fig. 1C. Initially, HBladFB-K4T fibroblasts were incorporated into IPC-50 collagen to form a collagen raft, as described in Section 2.5. Subsequently, HBladEC-K4T epithelial cells were seeded onto the surface of the collagen raft and cultured at the air-liquid interface. 3D-UBMM formed 9 days after seeding the HBladEC-K4T epithelial cells onto the surface of the collagen raft.

**Fig. 1.**
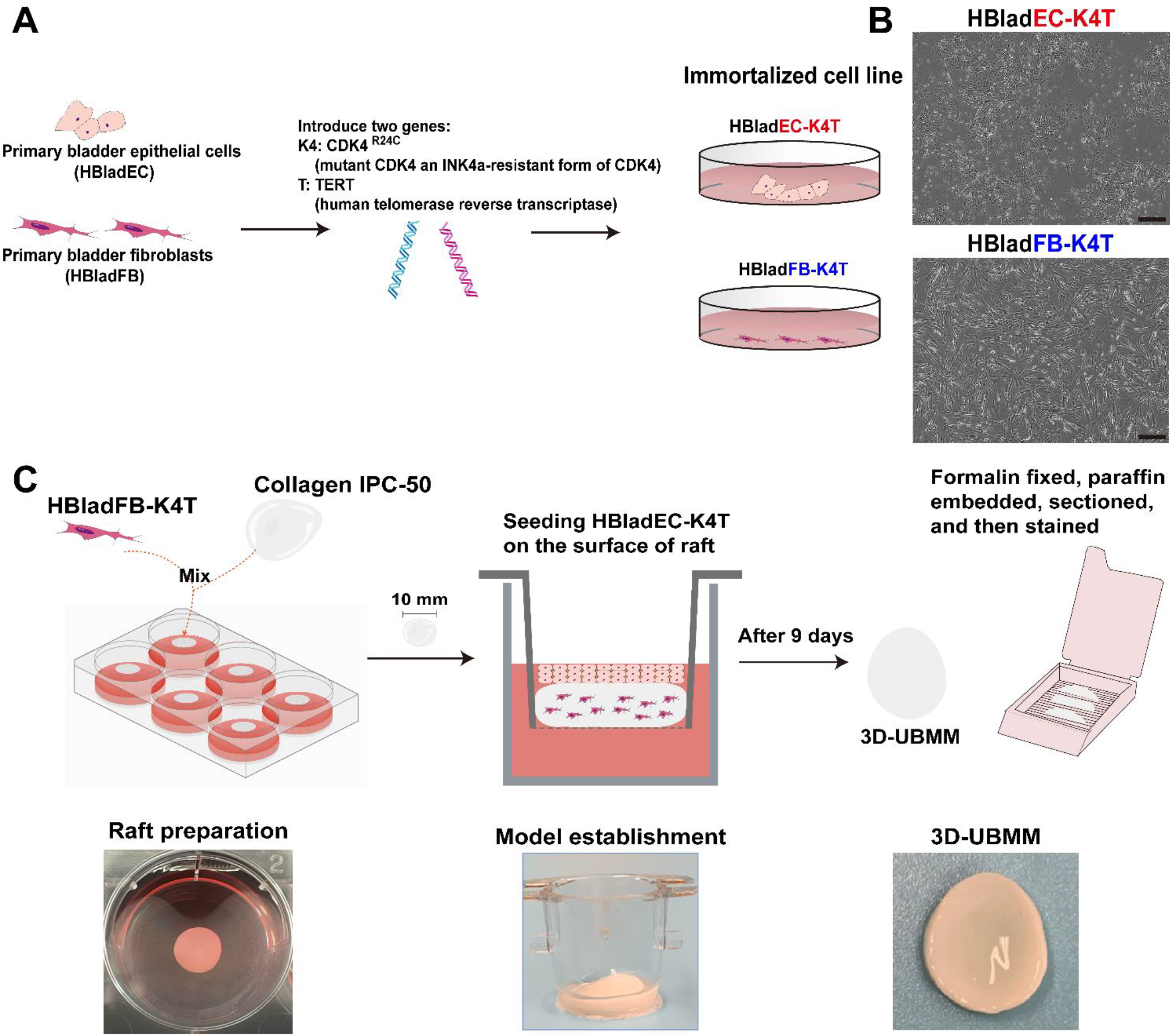
Establishment of immortalized primary HBladEC and HBladFB cell lines, and construction of the 3D-UBMM. A: Schematic illustration of immortalization of primary urothelial cells and fibroblasts by introducing genes TERT and CDK4^R24C^; B: Representative photos of immortalized HBladEC-K4T and HBladFB-K4T cells. The scale bar indicates 100 μm.; C: Construction of the 3D-UBMM.

### 3.2. Characteristics of the 3D-UBMM

As shown in Fig. 2A, the mucosal layers of the human urinary bladder are composed of upper urothelial cells and underlying connective tissue with fibroblast cells, forming the mucosa and the submucosa structures. Similarly, the upper layer of the 3D-UBMM consists of HBladEC-K4T epithelial cells attached to a collagen raft, which is suffused with HBladFB-K4T fibroblasts, resembling the mucosal and subepithelial structures of the human urinary bladder.

**Fig. 2.**
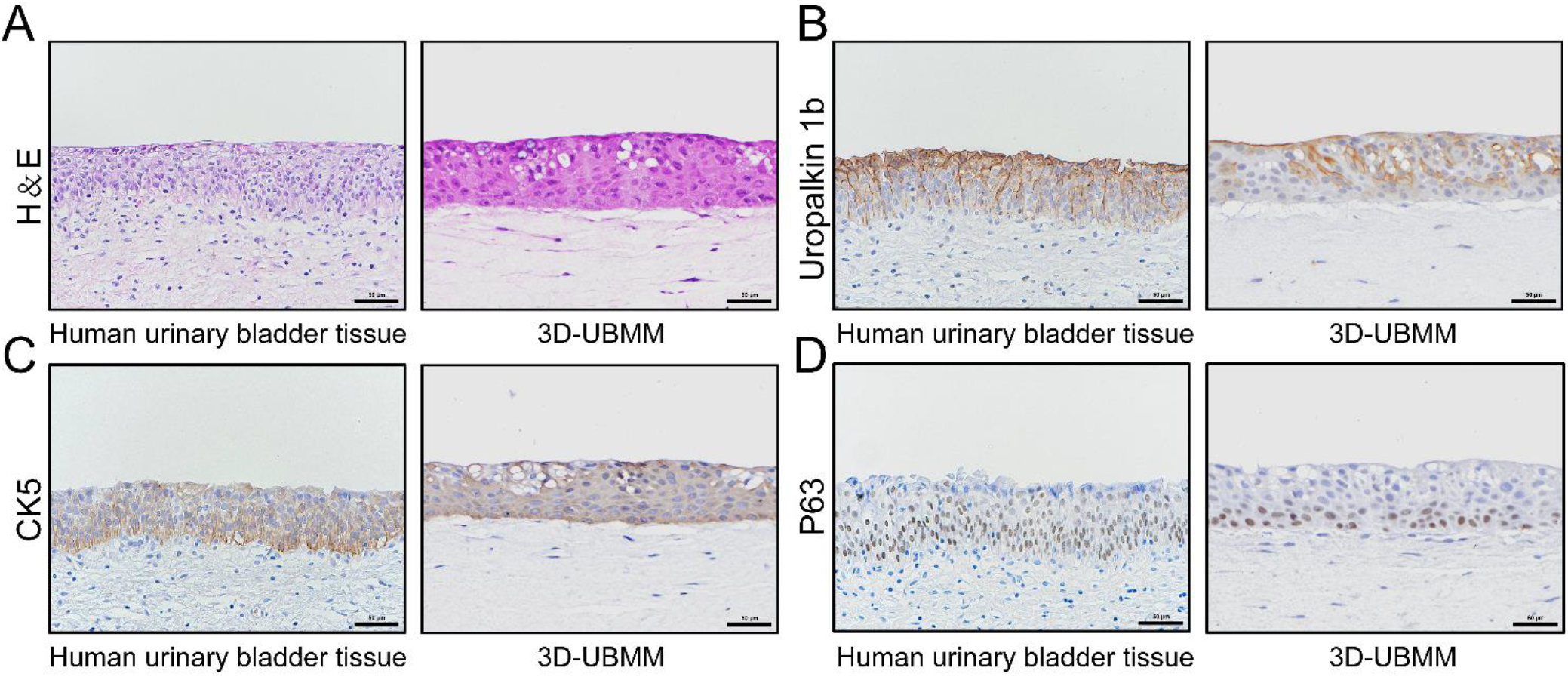
Characteristics of 3D-UBMM. A: the morphology of human urinary bladder epithelium and the 3D-UBMM; B-D: the expression of three marker proteins, UPK1B, CK5, and P63 in human urinary bladder epithelial cells and the 3D-UBMM. Representative images were acquired using an optical microscope. The scale bar indicates 50 μm.

The urinary bladder epithelium is composed of three cellular layers: an apical umbrella cell layer, an intermediate layer that can be several cells thick, and a basal stem cell layer. As shown in Fig. 2B-D, UPK1B, a specific marker for urothelial cells, is positive in the membrane and cytoplasm of urothelial cells in the apical umbrella cell layer and cells in the middle layers but negative in the basal layer of the 3D-UBMM. The expression pattern of UPK1B is very similar to the normal human urinary bladder epithelium. The expression pattern of CK5 in the 3D-UBMM is also similar to the normal human urinary bladder epithelium: the urothelial cells in the basal layer of the human bladder epithelium are positive for CK5, while some urothelial cells in the middle layer and a few urothelial cells in the apical umbrella cell layer are also positive. P63, a stem cell marker for the basal cells of the human urinary bladder epithelium, is expressed in the nucleus, and P63-positive cells are predominantly localized in the basal layer of the 3D-UBMM epithelium. Similarly, the urothelial cells in the basal layer of the human bladder epithelium are positive for P63, while some urothelial cells in the middle layer are also positive.

These findings show that the 3D-UBMM and human urinary bladder epithelium have similar histological and biological features.

### 3.3. Evaluation of the cytotoxicities of iAs^III^ and DMA^V^ in 2D-culture HBladEC-K4T

Fig. 3 illustrates the viability of HBladEC-K4T epithelial cells treated with iAs^III^ and DMA^V^ in 2D culture. The viability of HBladEC-K4T cells diminished in correlation with increasing concentrations of iAs^III^ and DMA^V^. The LC_20_ and LC_50_ value of iAs^III^ for HBladEC-K4T epithelial cells was 4.0 μM and 9.5 μM, respectively. The LC_20_ and LC_50_ values of DMA^V^ for HBladEC-K4T epithelial cells were 0.5 mM and 1.9 mM, respectively.

**Fig. 3.**
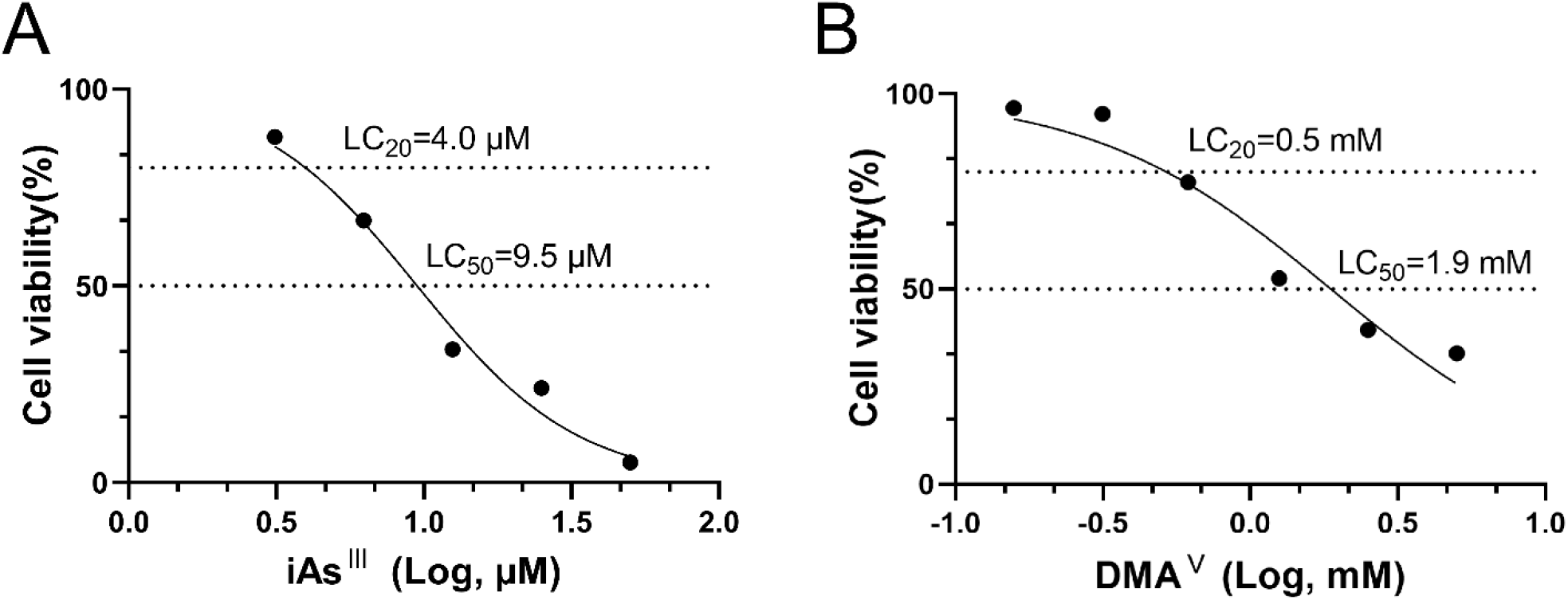
Evaluation of the cytotoxicity of iAs^III^ (A) and DMA^V^ (B) in 2D-cultures of HBladEC-K4T epithelial cells. The LC_20_ and LC_50_ values were calculated by nonlinear regression analysis using GraphPad prism 8. The LC_20_ and LC_50_ of iAs^III^ in HBladEC-K4T epithelial cells were 4.0 μM and 9.5 μM. The LC_20_ and LC_50_ of DMA^V^ in HBladEC-K4T epithelial cells were 0.5 mM and 1.9 mM.

### 3.4. Evaluation of the cytotoxicities of iAs^III^ and DMA^V^ using the 3D-UBMM

3D models more closely resemble the *in vivo* microenvironment, and previous studies have reported a noticeable tolerance to drugs in 3D cell models of tumor cells (Baru et al., 2022; Nowacka et al., 2022). Based on these findings, we selected concentrations of iAs^III^ and DMA^V^ exceeding the LC_50_ values identified in the 2D models using HBladEC-K4T epithelial cells to evaluate their toxicity in the 3D-UBMM. First, we evaluated the cytotoxicity of 40 μM iAs^III^ and 4 mM DMA^V^ in the 3D-UBMM. However, these levels of iAs^III^ and DMA^V^ resulted in the death of all of the urothelial cells in the 3D-UBMM. Therefore, we evaluated the cytotoxicity of 20 μM iAs^III^ and 2 mM DMA^V^. Apparent increases in necrotic cells were observed in the 3D-UBMM treated with 20 μM iAs ^III^ and 2 mM DMA^V^ compared to the control (Fig. 4A). P63-positive urothelial cells were decreased in the 3D-UBMM treated with 20 μM iAs^III^ and 2 mM DMA^V^ compared to the control (Fig. 4B). Furthermore, γ-H2AX-positive cells were readily apparent in the urothelial cells of 3D-UBMM treated with 20 μM iAs^III^ and 2 mM DMA^V^, whereas no γ-H2AX-positive urothelial cells were observed in the controls (Fig. 4C). This indicated that iAs^III^ and DMA^V^ induced DNA damage in urothelial cells in the 3D-UBMM.

**Fig. 4.**
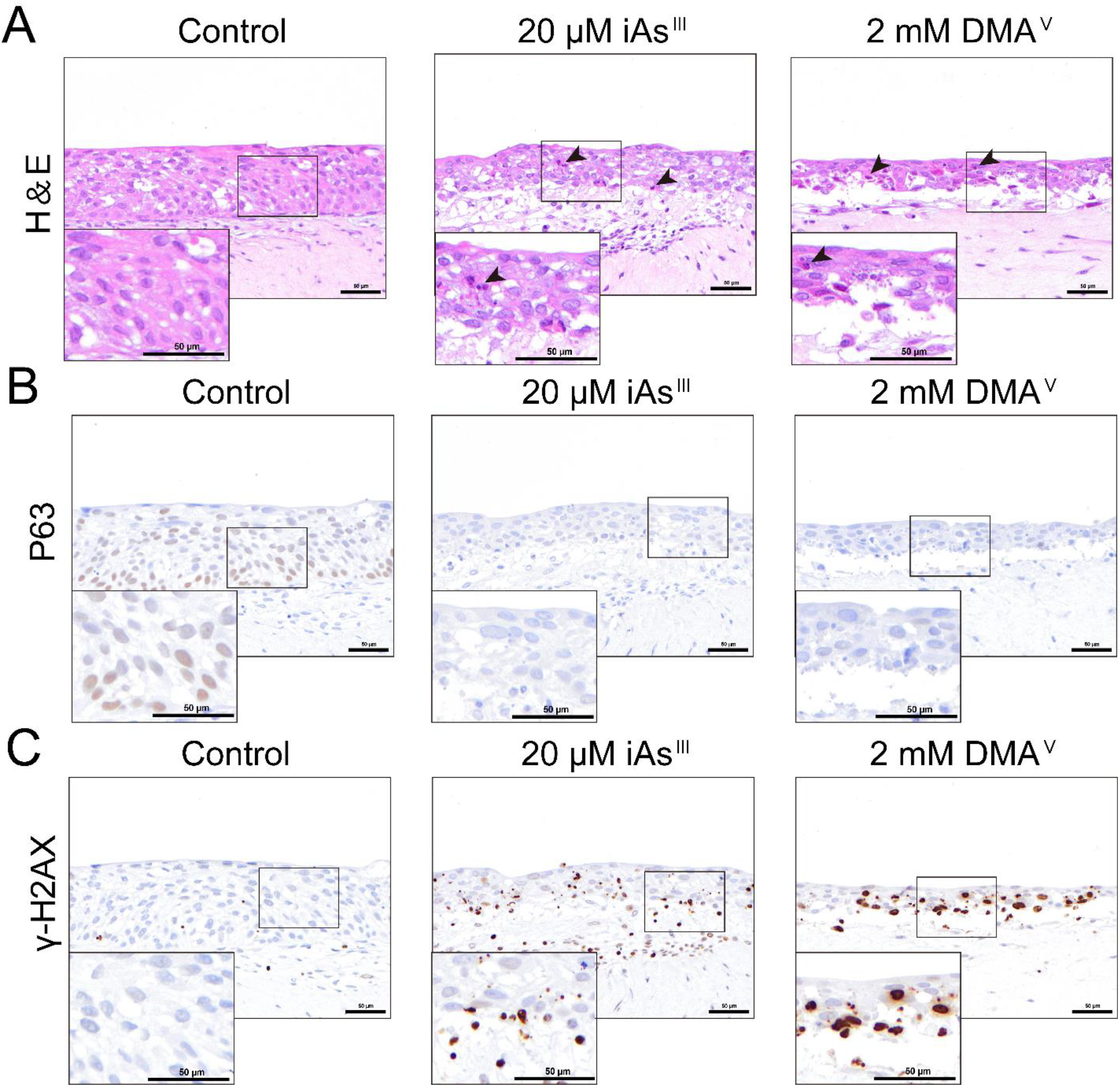
Evaluation of the cytotoxicity of 20 μM iAs^III^ and 2 mM DMA^V^ using the 3D-UBMM. A: Representative photos of histological alterations of 3D-UBMM treated with 20 μM iAs^III^ and 2 mM DMA^V^. Black arrows point to necrotic cells. B-C: Expression of P63 and γ-H2AX in 3D-UBMM treated with 20 μM iAs^III^ and 2 mM DMA^V^. The scale bar indicates 50 μm.

To determine the dose-response relationship of the cytotoxicity of iAs^III^ and DMA^V^, 3D-UBMM were treated with iAs^III^ at doses of 5 μM, 10 μM, and 20 μM, and DMA^V^ at doses of 0.5 mM, 1 mM, and 2 mM. The lowest dosages were set at approximately the LC_20_ of iAs^III^ and DMA^V^. 5 μM iAs^III^ and 0.5 mM DMA^V^ did not cause histopathologic alterations compared with the control (Fig. 5A). Neither necrotic cells nor γ-H2AX-positive cells were observed in the 3D-UBMM following treatment with 5 μM iAs^III^ and 0.5 mM DMA^V^ (Fig. 5A and 5C). Additionally, there was no significant difference in the number of P63-positive cells between the control and the treated 3D-UBMM, indicating an absence of toxicity in the 3D-UBMM treated with 5 μM iAs^III^ and 0.5 mM DMA^V^ (Fig. 5B). With increasing concentrations of iAs^III^ and DMA^V^, the urothelial cells in the 3D-UBMM displayed a marked rise in necrosis. P63-positive basal cells were decreased in a dose-dependent manner following iAs^III^ and DMA^V^ treatment (Fig. 5B and 5D). iAs^III^ significantly decreased the number of P63-positive cells at 10 and 20 μM compared to controls, while DMA^V^ caused a significant reduction at 2 mM. Furthermore, iAs^III^ and DMA^V^ increased the number of γ-H2AX-positive cells in a dose-dependent manner, with significant increases observed at 20 μM iAs^III^ and 2 mM DMA^V^ compared to the controls (Fig. 5C and 5E).

**Fig. 5.**
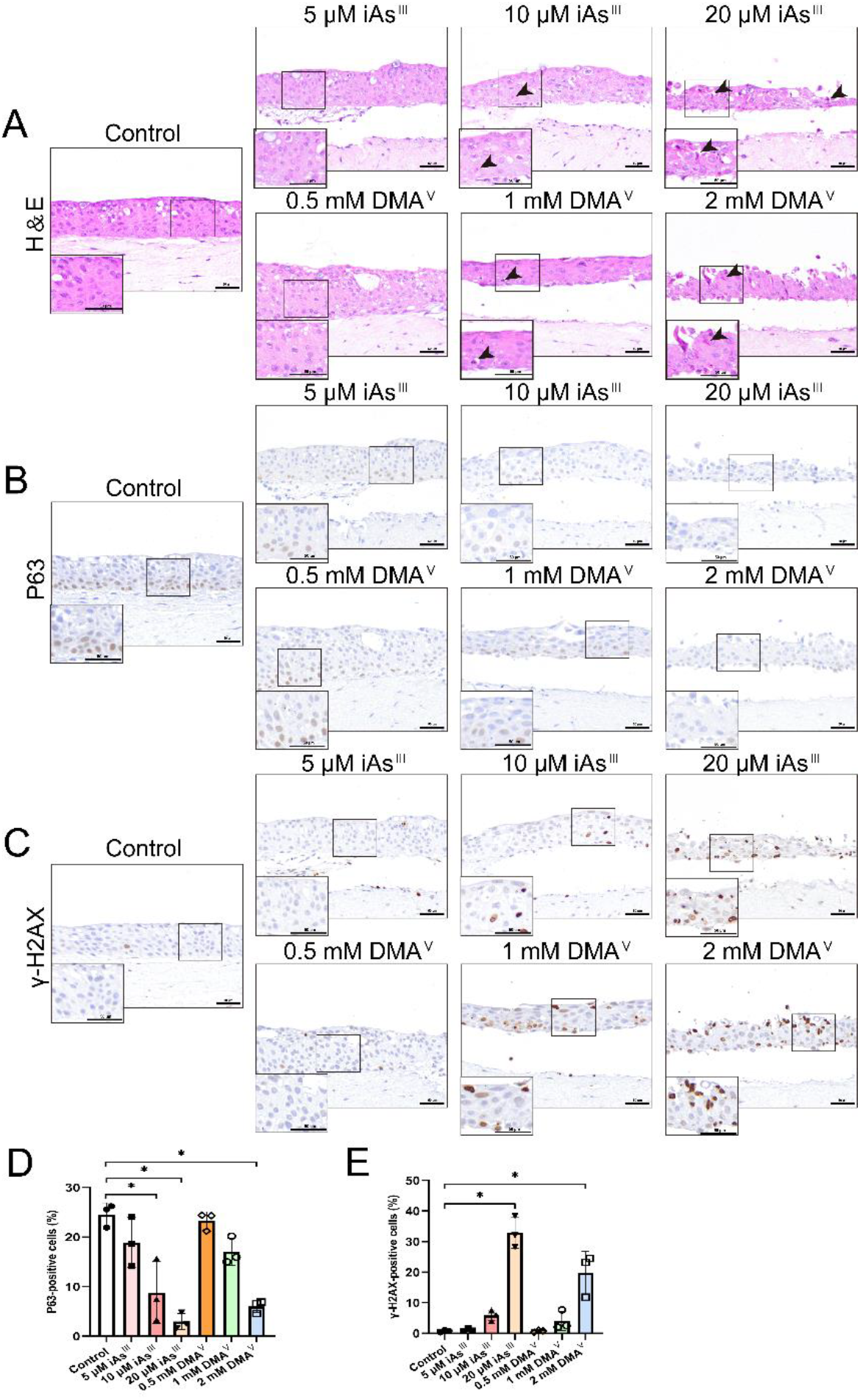
Cytotoxicity of increasing doses of iAs^III^ and DMA^V^. A: Representative photos of histological alterations. Black arrows point to necrotic cells. B-C: Representative images of P63 and γ-H2AX expression in 3D-UBMM exposed to three concentrations of iAs^III^ and DMA^V^. A high-magnification view is shown in the lower-left corner of each image, corresponding to the area outlined by a small box in the center image. The scale bar indicates 50 μm. D: Percentage of P63-positive cells. iAs^III^ significantly decreased the number of P63-positive cells at 10 and 20 μM compared to the controls. *p < 0.01. DMA^V^ significantly decreased the number of P63-positive cells at 2 mM compared to the controls. *p < 0.01. E: Percentage of γ-H2AX-positive cells. iAs^III^ and DMA^V^ significantly increased the number of γ-H2AX-positive cells at 20 μM and 2 mM compared to the controls, respectively. *p < 0.01. Note that the photo of p63 in the 3D-UBMM control is also shown in Figure 2.

## 4. Discussion

Arsenic is a well-established human bladder carcinogen, with cytotoxicity playing a pivotal role in its carcinogenicity. However, a comprehensive understanding of the cytotoxic effects of arsenic on the human urinary bladder epithelium remains elusive. Therefore, we developed a novel 3D urinary bladder mucosa model (3D-UBMM) using immortalized human bladder fibroblast cells and urothelial cells. The applicability of our 3D-UBMM was tested by evaluating the cytotoxicity of two *in vivo* carcinogens, a stronger carcinogen iAs^III^ and a weaker carcinogen DMA^V^.

The human urinary bladder mucosal layer consists of an overlying urothelium composed of three layers of specialized epithelial cells and a lower connective tissue layer. Our 3D-UBMM also has a bilayer structure with an upper layer of human urothelial cells and a lower layer composed of a collagen raft containing human urinary bladder fibroblast cells. The epithelial cell layers of the urothelium consist of apical umbrella cells, an intermediate layer of epithelial cells, and a basal stem cell layer. UPK1B, CK5, and P63 were selected to evaluate the histology of the 3D-UBMM. UPK1B is expressed almost exclusively in the urinary bladder epithelium(Dalghi et al., 2020). UPK1B is normally expressed in apical umbrella cells and intermediate cells of the urothelium. CK5, a member of the cytoskeletal protein family exhibits high expression in the basal layer of the urothelium and lower expression in the intermediate layer (Wiessner et al., 2022). It contributes to the formation of intermediate filaments, thereby supporting cellular structural stability and morphology (Al-Sharaky et al., 2021). P63 is a basal layer cell marker within the human urinary bladder epithelium. P63-expressing cells are situated at the base of the bladder epithelium and adhere firmly to the connective tissue layer, playing an essential role in the structural integrity and regenerative capacity of the urinary bladder epithelium (Pignon et al., 2013). As can be seen from the results of the present study, the 3D-UBMM closely resembles the human urinary bladder epithelium in terms of morphology and expression of the marker proteins UPK1B, CK5, and P63.

Although arsenicals do not directly interact with DNA, and consequently, are not direct-acting genotoxicants, oxidative stress has been proposed as an indirect mechanism contributing to their genotoxic potential (Medda et al., 2021; Vachiraarunwong et al., 2024; Wei et al., 2002). Reactive oxygen species (ROS) have been reported to be generated during arsenic metabolism in the cell and can cause oxidative damage to DNA, proteins, and lipids (Cohen et al., 2013; Vachiraarunwong et al., 2024). Excessive ROS generation can exacerbate cellular injury, creating a feedback loop that amplifies both cytotoxic and genotoxic stress (Cohen et al., 2013; Vachiraarunwong et al., 2024). Although we did not directly assess ROS production in this study, excessive ROS generation has been implicated in cytotoxic and genotoxic mechanisms in other systems (Cohen et al., 2013; Vachiraarunwong et al., 2024). In our study, administration of iAs^III^ and DMA^V^ resulted in a dose-dependent increase in γ-H2AX-positive cells, a well-recognized marker of DNA double-strand breaks. While γ-H2AX does not indicate direct genotoxicity, it serves as a sensitive biomarker of DNA damage potentially resulting from sustained oxidative stress or other indirect mechanisms (Suzuki et al., 2020; Toyoda et al., 2015). We also observed a dose-dependent decrease in P63-positive cells in the 3D-UBMM. Given that basal cells function as a stem cell reservoir in the bladder urothelium, their depletion may compromise epithelial integrity and regenerative capacity, potentially contributing to carcinogenic progression over time.

It is now widely recognized that 3D cell models may more accurately reflect the *in vivo* situation compared to 2D cell cultures. In a study by Carrie J. Lovitt, it was demonstrated that breast cancer cells cultured in a 3D *in vitro* model can form an extracellular matrix, which is closely related to both drug delivery and tumor cell growth. In contrast, breast cancer cells in 2D culture struggled to establish an extracellular matrix (Lovitt et al., 2018). Marta Nowacka’s research also showed that the 3D ovarian cancer cell model exhibited greater drug resistance compared to ovarian cancer cells in 2D culture (Nowacka et al., 2022). In our study, no necrotic cells or γ-H2AX-positive cells were observed in the 3D-UBMM treated with 5 μM iAs^III^ or 0.5 mM DMA^V^, while the same concentrations in 2D-cultured HBladEC-K4T resulted in about 20% cell death. Importantly, during proliferation and cell differentiation of basal layer stem cells, some cells migrate upwards and differentiate into umbrella cells, forming the apical layer of the urinary bladder epithelium. This umbrella structure plays an important role in protecting the urinary bladder epithelium from irritation and chemicals in the urine (Dalghi et al., 2020; Jafari and Rohn, 2022). While this structure is difficult to organize in 2D cell cultures of bladder urothelial cells, the 3D-UBMM does allow for the formation of an apical umbrella structure. Therefore, the tolerance of the 3D-UBMM to iAs^III^ and 0.5 mM DMA^V^ is higher than that of 2D-cultures.

Urinary concentrations of iAs^III^ and DMA^V^ in individuals living in arsenic-contaminated areas have been reported to range up to approximately 0.26 μM and 1.1 μM, respectively (Hata et al., 2012).

Although the highest concentrations of iAs^III^ (20 μM) and DMA^V^ (2 mM) used in this study exceed urinary concentrations observed in exposed human populations, they were selected to enable mechanistic evaluation of cytotoxic responses in the 3D-UBMM model. Given the limitations in translating *in vitro* exposures to *in vivo* human conditions, particularly due to differences in exposure duration and bioavailability, these concentrations were used to define toxic thresholds and explore potential cellular responses. Future studies incorporating time-course analyses and comprehensive gene expression profiling will be necessary to better correlate *in vitro* findings with physiologically relevant exposure levels.

In conclusion, we have constructed a novel 3D-UBMM with characteristics similar to human urinary bladder epithelium. The cytotoxicity and DNA damage induced by iAs^III^ and DMA^V^ were tested in the 3D-UBMM and reflected the stronger cytotoxicity of iAs^III^ compared to DMA^V^ in mice and humans. Suzuki et al. and Tokar et al. (Suzuki et al., 2008; Tokar et al., 2011)reported that mice exposed to Sodium arsenite (iAs^III^) developed urinary bladder hyperplasia while Arnold et al. (Arnold et al., 2006) reported that no hyperplastic effects, or tumors, were observed in mice exposed to DMA^V^. In humans, there is sufficient evidence that iAs^III^ is a urinary bladder carcinogen, while DMA^V^ is classified as possibly carcinogenic in humans, but this classification is based on animal studies and no studies on the effect of DMA^V^ in humans are presented (IARC, 2012). Thus, the IARC report strongly suggests that DMA^V^ is less carcinogenic in humans than iAs^III^. Importantly, while the cytotoxicity of iAs^III^ and DMA^V^ in 2D-cultures of HBladEC-K4T cells differed by approximately 125 to 200-fold, exposure of 3D-UBMM to approximately 2D-culture LD50 levels of these arsenicals induced significant formation of γ-H2AX, suggesting that their mode of action was similar. These findings indicate the effectiveness of the 3D-UBMM in evaluating arsenic toxicity. Importantly, one of the inconsistencies in the *in vivo* data is that while iAs^V^ and iAs^III^ are known human urinary bladder carcinogens and also induce urinary bladder cancer in rats, there are no studies demonstrating that iAs^V^ or iAs^III^ induce urinary bladder cancer in mice. Future studies examining the time-dependent response and comprehensive gene expression profiling of arsenicals and other bladder toxins using the 3D urinary bladder mucosa model are warranted to further elucidate underlying mechanisms of toxicity and carcinogenicity, and may provide information helping to resolve the inconsistencies noted above and increase our understanding of bladder toxins. In addition, the 3D-UBMM can be used to investigate the carcinogenicity of DMA^V^. DMA^V^ is metabolized differently by rats than by mice and humans, and while it is a rat urinary bladder carcinogen it does not induced urinary bladder cancer in mice and its effect in humans is currently unknown.

## Acknowledgments

The authors gratefully acknowledge the technical assistance of Rie Onodera, Keiko Sakata, Yuko Hisabayashi, and Yukiko Iura (Department of Molecular Pathology, Graduate School of Medicine School, Osaka Metropolitan University Osaka, Japan).

